# Genome-wide CRISPR Screening Reveals Cullin-1 as a Therapeutic Target Enhancing Efficacy of CD19-directed Immunotherapy

**DOI:** 10.64898/2026.07.26.740751

**Authors:** Narendra V Gottumukkala, Tomáš Loja, Michal Šmída

**Affiliations:** Central European Institute of Technology (CEITEC), Masaryk University, Brno, Czech Republic; Department of Internal Medicine-Hematology and Oncology, University Hospital Brno and Faculty of Medicine, Masaryk University, Brno, Czech Republic

**Keywords:** CD19, Cullin-1, Pevonedistat, Cancer immunotherapy, Antigen escape, CAR-T cells, CD81

## Abstract

Cancer immunotherapy targeting B-cell specific CD19 antigen meant a major breakthrough in the treatment of B-cell malignancies. Yet, vast proportion of treated patients experience relapse and failure of the therapy. Although multiple mechanisms of CD19-immunotherapy failure have been described, CD19-negative relapses represent the major hurdle in achieving higher and durable response rates.

Our established *in vitro* co-culture models revealed that suboptimal CAR-T cell performance, inefficient to mediate target cell killing, results in robust downregulation of CD19 target antigen. Using genome-wide CRISPR screening, we addressed the mechanisms responsible for such CD19 downregulation and identified Cullin-1 and CD81 playing instrumental role in negative and positive regulation of CD19 expression, respectively. Inhibiting Cullin-1 activity with pevonedistat prevents the loss of CD19 under immunotherapeutic pressure, results in higher CD19 surface levels and consequently enhances the efficacy of target cell killing by CD19-CAR-T cells, CD19-CAR-NK cells and CD19-targeting antibody treatment. Mechanistically, we show that pevonedistat blocks the degradation of CD19 upon its internalization and allows its recycling back to the plasma membrane. CD81 chaperone protein is critically involved in this process as the absence of CD81 abrogates the effect of pevonedistat.

In summary, we identify Cullin-1 as a novel and druggable regulator of CD19 protein stability. Cullin-1 inhibition augments CD19 surface expression, thereby improving the efficiency of CD19-targeting immunotherapies, thus arguing for potential incorporation of pevonedistat into novel combination therapies.

**Key Points:**

- Cullin-1 inhibition stabilizes CD19 surface expression, preventing its loss under immunotherapeutic pressure
- Pevonedistat treatment enhances the efficacy of CD19-CAR-T cells, CAR-NK cells and CD19-targeting antibodies

## Introduction

Chimeric Antigen Receptor (CAR) T cell therapy directed against CD19 has transformed the treatment of relapsed or refractory B cell malignancies, including acute lymphoblastic leukemia (ALL) and Non-Hodgkin lymphomas (NHL), leading to the approval of multiple CAR-T products^1–3^. Despite these remarkable advances, a significant fraction of patients experiences disease relapse, most commonly due to the loss or downregulation of CD19 antigen expression on the tumor cell surface^4,5^. Antigen escape represents a major biological barrier to durable remissions, making the identification of strategies to restore and stabilize CD19 an urgent clinical priority.

Several distinct mechanisms have been described so far that can contribute to treatment failure, such as acquired *CD19* genetic mutations, alternative splicing of the CD19 locus, hypermethylation-mediated silencing of *CD19* promotor or cell lineage switch^6–10^. The majority of resistant cases, however, retain intact CD19 mRNA transcripts but display drastically diminished surface protein density^4,5^. This discrepancy strongly indicates that post-transcriptional or post-translational mechanisms such as altered intracellular trafficking, membrane retention, or accelerated protein degradation can drive antigen loss under immunotherapeutic selective pressure. However, the precise molecular pathways regulating CD19 protein stability at the plasma membrane remain poorly defined.

The functional presentation of CD19 at the cell surface is highly dependent on its interaction with chaperone proteins. CD19 relies heavily on the tetraspanin chaperone CD81 for proper export from the endoplasmic reticulum, stable surface expression, and organization into membrane microdomains^11^. These microdomains are critical during immunotherapy, as effector cell cytotoxicity, whether driven by CAR-T, CAR-NK, or monoclonal antibodies, requires sustained, high-density antigen engagement to form a productive immunological synapse^12^. When surface CD19 is internalized and degraded rather than recycled, these antigen levels drop below the critical threshold required to trigger robust effector cell degranulation and tumor cell lysis.

Because intracellular protein turnover is primarily governed by ubiquitination, E3 ubiquitin ligases represent highly logical potential suspects that might be involved in the rapid downregulation of surface receptors. Cullin-RING ligases (CRLs) comprise the largest family of E3 ubiquitin ligases, mediating the ubiquitination and subsequent degradation of numerous cellular substrates^13^. CRL1, which is built upon a Cullin-1 (CUL1) scaffold, requires activation via covalent attachment of the ubiquitin-like protein NEDD8. Pharmacologic inhibition of CRL activity can be achieved using pevonedistat (MLN4924), a first-in-class small molecule inhibitor of the NEDD8-activating enzyme (NAE) that is currently under clinical investigation for hematologic malignancies and solid tumors^14,15^.

Here, we have established *in vitro* cell models of suboptimal CAR-T performance, which results in robust CD19 surface antigen downregulation. Using an unbiased, genome-wide CRISPR knockout screen under CAR-T immune pressure, we identified two key genetic regulators of CD19 antigen dynamics: the ubiquitin ligase scaffold CUL1 and the tetraspanin chaperone CD81. We demonstrate that pharmacologic inhibition of CRL1 with pevonedistat restores CD19 surface levels across multiple lymphoma and leukemia models. Mechanistically, CUL1 inhibition by pevonedistat does not block antigen internalization; rather, it coordinates CD81 upregulation and accelerates CD81-dependent surface recycling. This rapid recycling stabilizes effector-target conjugates and maintains the high antigen density required for robust immune synapse formation. Consequently, pevonedistat synergistically re-sensitizes malignant B cells and primary patient samples to a diverse array of immunotherapeutic modalities, including CAR-T, CAR-NK, and antibody-dependent cellular cytotoxicity (ADCC). These findings establish CRL inhibition as a promising, readily translatable therapeutic strategy to minimize CD19-negative antigen escape.

## Material and Methods

### Cell cultures

Burkitt lymphoma cell lines Ramos and Raji, and HeLa cell line were obtained from ATCC, chronic lymphocytic leukemia (CLL) cell line MEC1, T-cell lines Jurkat and HSB-2 were obtained from DSMZ and CLL cell line HG3 was kindly provided by the lab of Dr. Rosenquist (Uppsala, Sweden). MEC1 cell line was cultured in IMDM medium, while all the others were cultured in RPMI 1640. Media were supplemented with 10% heat-inactivated fetal bovine serum (FBS) and 1% penicillin/streptomycin (all from Sigma-Aldrich).

NK cell lines NK92MI and KHYG16 were kindly provided by Dr. Milan Vrabel (IOCB, Prague) and cultured in complete RPMI supplemented with 10% FBS, 10% Horse serum, 1% pen strep and 100 U/mL IL-2. Primary CLL samples were selected from the biobank of the Department of Internal Medicine-Hematology and Oncology (University Hospital Brno) and cultured in AIM-V medium (Sigma-Aldrich) supplemented with 10% FBS, 1% penicillin/streptomycin, 100 U/ml IL-2, 100 ng/ml Resiquimod and 50 mM 2-ME.

All cell cultures were maintained at 37 °C in 5% CO_2_ atmosphere.

### Generation of CD19-CAR-T cells

Lentiviral particles were generated using the CD19-targeting CAR construct containing a 4-1-BB costimulatory domain (Addgene #135992). Lenti-X cells (Clontech) were seeded in T150 flasks at 60% confluency and transfected next day with equimolar ratios of pMD2.G (#12259), psPAX2 (#12260) and pSLCAR-CD19 using polyethylenimine (PEI) at a 3:1 ratio (PEI:DNA) in reduced serum Opti-MEM (ThermoFisher). The medium was replaced 3 hours post-transfection. Viral supernatants were harvested at 48 and 72 hours, clarified, and concentrated using Lenti-X Concentrator (TakaraBio) according to the manufacturer’s protocol. The resulting viral pellets were resuspended in AIM-V medium.

Buffy coats from healthy donors were purchased from the University Hospital Brno. T cells were isolated using Ficoll gradient centrifugation and RosetteSep T cell enrichment kit (StemCell). T cells were activated by culturing with CD3/CD28 magnetic beads (ThermoFisher) at 1:3 beads:cell ratio in AIM-V medium for 24h and transduced using a retronectin-based spinfection protocol. Briefly, 1.5 mL microcentrifuge tubes were coated with 100 µg/ml Retronectin (TakaraBio) for 2h at room temperature (RT), followed by blocking with 2% BSA/PBS. T cells (1×10^6^) were resuspended in 200 µL of concentrated lentivirus with 10 μg/ml Polybrene (Merck) in the pre-coated tubes and centrifuged at 1500 g for 2hours at 32°C. Cells were incubated for 6 hours before a complete media exchange and CD19-CAR expression was evaluated 72 hours post-transduction by surface staining with AlexaFluor-647 conjugated FMC63 scFv monoclonal antibody (Custoscan). Cells were expanded in AIM-V medium supplemented with 100 U/ml IL-2.

### Genome-wide CRISPR screening

Human genome-wide sgRNA knockout library Brunello was obtained from Addgene (#73179)^16^. Lenti-X cells were seeded in four T150 flasks at 50-60% confluency and next day transfected with equimolar ratios of pMD2.G, psPAX2 and Brunello plasmid using polyethylenimine (PEI) at a 3:1 ratio (PEI:DNA) in reduced serum Opti-MEM (ThermoFisher). Medium was replaced 3h post-transfection and viral supernatants harvested at 48h and 72h, clarified, and concentrated 50-fold using Lenti-X Concentrator into complete DMEM.

Ramos cells (1×10^8^/replicate) were transduced in two replicates with the predetermined viral library titer ensuring multiplicity of infection (MOI) of approximately 0.3 and cells were selected with puromycin. These cells were cultured for 14 days and then mixed with primary control T cells or CAR-T cells at an E:T ratio of 1:4 for 48hours. Cultures were stained with CD19-APC and sorted by FACS into CD19-high and CD19-low cell population. Genomic DNA was extracted and sgRNA cassettes PCR amplified using NEBNext Ultra II Q5 (NEB) with a mix of 8 staggered P5 primers and a sample-specific indexed P7 primer (**Suppl. Table 1**). Libraries were purified on AMPure XP beads (BeckmanCoulter) and sequenced with the Nextseq 500/550 High output v2 kit on Illumina NextSeq 500 sequencer using 75 cycles. Sequencing data were quality checked using FastQC^17^ and adaptors and low-quality ends were trimmed using trimmomatic tool^18^. Sequencing reads were aligned to the sgRNA library file and read counts were normalized. Fold change, p-values and false discovery rates (FDR) were calculated for enrichment or depletion of each sgRNA within the sorted population compared to their abundance in control T cell-treated cultures. The analysis was done using freely available softwares MAGeCK and EdgeR^19,20^ and gene ontology enrichment analysis was performed using online tool Gorilla^21^.

### Generation of knockout cell lines

Individual sgRNAs targeting *CUL1* (CCGGGTTGACGACATTGTGA) and *CD81* (CATGATCACAGCGACCACGA) genes were cloned into lentiGuide-mCherry plasmid, lentiviral particles produced and Ramos B cells stably expressing Cas9 transduced. mCherry-positive population was sorted to generate CUL1 knockouts. For CD81-knockout cells, cells were stained with CD81-APC and CD81-negative fraction was sorted.

### Effector : Target cell co-culture assays

Target B cells were stably transduced with Azurite plasmid (Addgene, #36086) for easy FACS identification. CAR-transduced effector cells co-expressed GFP. Effector cells were mixed with target cells at predetermined effector to target (E:T) ratios and incubated for specified time intervals in U-bottom 96-well-microplates (TPP). For ADCC, CD19 antibody Tafasitamab (10,6 mg/ml, Incyte Biosciences) was added together with CD16-expressing KHYG cells. In some experiments, pevonedistat (10 nM), Dynasore (1 µM), bafilomycin A (100 nM), chloroquine (100 nM), and MG132 (100 nM) (all from MedChemExpress) were added simultaneously with the initiation of the co-culture.

CD19 expression was determined by flow cytometry measurement of CD19-APC staining, gating on azurite-positive cells. Cell killing was analyzed via flow cytometry by measuring the remaining viable azurite-positive cells through 7-AAD viability dye exclusion and normalizing these numbers to the viability in control effector cell cultures.

### Flow cytometry

Cells were washed with PBS and incubated with respective antibodies (**Suppl. Table 2**) for 30 minutes at 4 °C. After PBS wash, they were resuspended in 300 µL PBS and immediately analyzed on FACSVerse (BD). Data analyses were performed using FlowJo software.

For conjugate formation, total events were gated to isolate doublet population based on FSC-A and FSC-H, which was subsequently evaluated on dot plot for both effector (CAR-GFP) and target (azurite) cell fluorescence. Double positive events were quantified as CAR:B cell conjugates.

### Quantitative real-time PCR

Total RNA was extracted with RNeasy Mini Kit (Qiagen) according to the manufacturer’s instructions. cDNA was prepared using RevertAid reverse transcriptase (ThermoFisher) and random hexamer primers (ThermoFisher). Quantitative PCR was conducted in triplicates using PowerUp SYBR Green MasterMix (Applied Biosystems) on a QuantStudio 12K Flex system (Applied Biosystems) with primers specific for *CD19* and *GAPDH*. Relative *CD19* expression was normalized against the housekeeping control *GAPDH*.

### Western blotting

Cells were lysed in RIPA buffer supplemented with PMSF and PhosSTOP (Roche) and boiled at 98°C for 10 min. Proteins were resolved by SDS-PAGE, transferred to nitrocellulose or PVDF membranes, and blocked in 5% BSA or 5% milk for one hour at RT. Membranes were probed overnight at 4°C with primary antibodies (**Suppl. Table 2**) and after 3x TBS/Tween20 washes, incubated for one hour at RT with HRP-conjugated secondary antibodies. Signals were developed using Clarity ECL (Bio-Rad) and captured on Alliance Q9 imaging system (Uvitec). To assess requirement of protein synthesis, cells were treated with 1 mg/mL cycloheximide (Sigma-Aldrich) together with pevonedistat for indicated times and lysed for western blot.

### HeLa cell assay

Stable CD19-positive HeLa cells were generated by transducing CD19 construct (Addgene, #201919) into HeLa azurite cells. They were seeded in 6-well-plate and effector cells (NK92MI-WT/CAR) were added next day at 1:5 ratio together with pevonedistat for 24hr. Target cell viability was assessed by quantifying the absolute number of surviving azurite-positive HeLa cells.

### Statistical analysis

All statistical analyses were performed using GraphPad Prism version 11.0.2. Data are presented as mean ± standard deviation (SD). For comparisons between two paired groups, statistical significance was determined using paired t-test. For experiments comparing three or more conditions, statistical significance was assessed using one-way analysis of variance (ANOVA), followed by multiple-comparisons testing. A p-value of less than 0.05 was considered statistically significant (*p <0.05, **p <0.01, ***p <0.001, ****p <0.0001).

## Results

### Suboptimal CAR T-cell and therapeutic CD19 antibody performance induces CD19 antigen downregulation

To investigate the consequences of suboptimal immunotherapeutic selective pressure, which is inefficient in killing target cells, we used primary CAR-T cells at a low effector-to-target ratio (E:T) to co-culture with CD19+ B-cell lymphoma and leukemia cell lines (Ramos, HG3, and Mec1). Upon this exposure, we observed a strong CD19 antigen loss on the malignant cells **(Figure 1A)**. To further model and explore this phenomenon *in vitro*, we established a co-culture of B-cell lines (Ramos, Mec1, HG3, and Raji) with Jurkat T cells expressing a CD19-targeting CAR construct. Interestingly, this Jurkat model resulted in a similarly profound loss of surface CD19 expression on all the malignant B-cell lines **(Figure 1B)**. This phenotype was highly reproducible in primary malignant cells, as evidenced by a similar downregulation of surface CD19 in primary cells obtained from seven patients with chronic lymphocytic leukemia (CLL) following Jurkat CAR-T exposure *ex vivo* **(Figure 1C)**. Finally, to address whether the loss of CD19 occurs only on the cell surface or whether total cellular CD19 protein is reduced, we performed a Western blot analysis, which confirmed a significant reduction in total CD19 protein levels across our models and also in two of the patient samples (Figure 1D, 1E).

**Figure 1.**
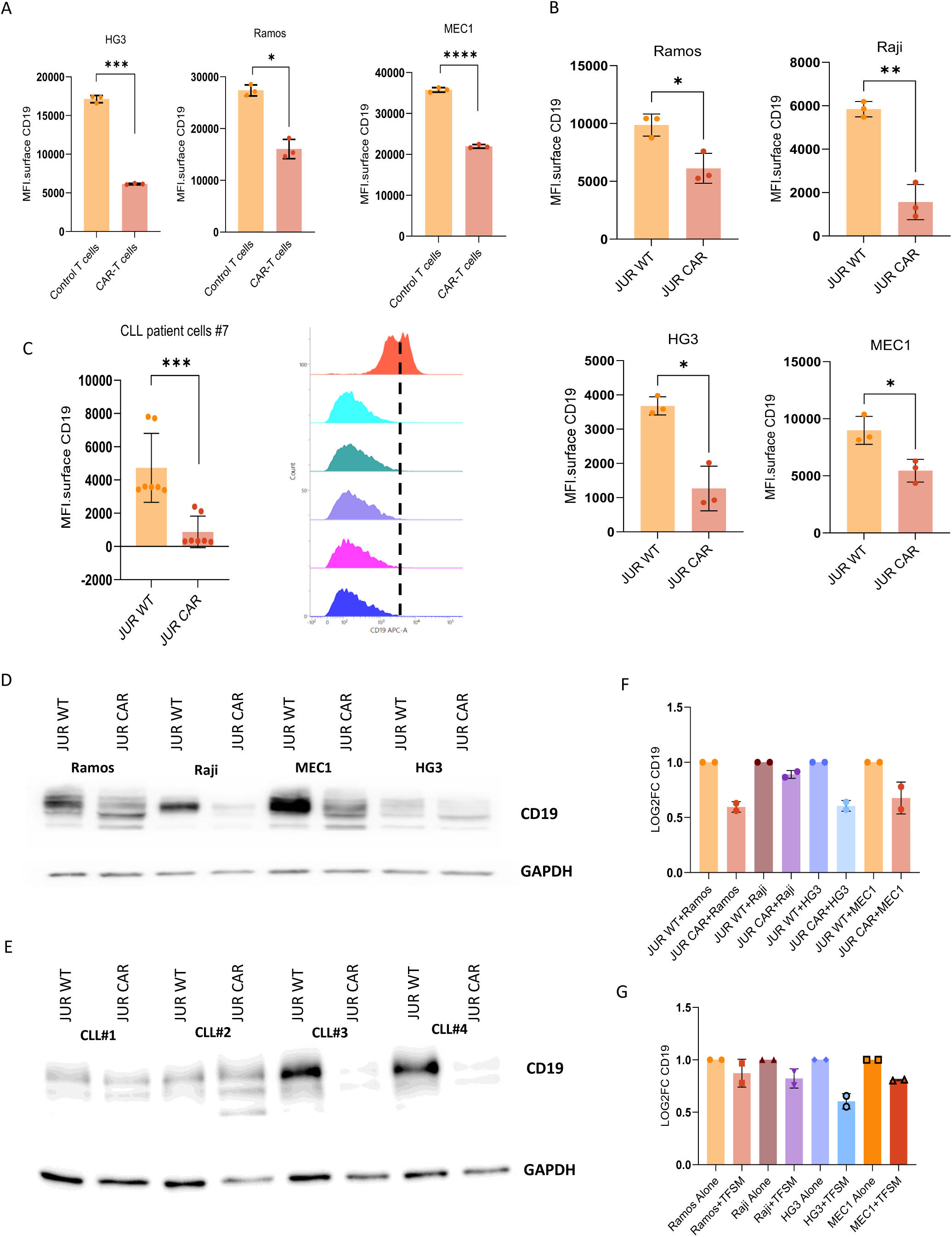
Suboptimal immunotherapeutic treatment results in CD19 antigen downregulation. (**A**) Ramos, HG3 and MEC1 B-cell lines were treated with primary human CD19-CAR-T cells or control T cells at a low effector : target (E:T) ratio of 1:3 for 48h. The expression levels of CD19 target antigen on the surface of B cells was determined by flow cytometry. Data represent mean of three independent biological replicates (n=3) ± SD. (**B**) Malignant B-cell lines Ramos, Raji, HG3 and MEC1 were cultured with control or CD19-CAR-expressing Jurkat T-cell line at 1:1 ratio for 48h and surface CD19 levels on the B cells were determined by flow cytometry; mean ± SD (n=3). (**C**) Primary patient CLL cells from 7 different patients were cultured with control or CD19-CAR-Jurkat cells for 48h at 1:1 ratio and CD19 surface levels were determined by flow cytometry. Left: mean ± SD (n=7). Right panel shows individual histograms for a sample cultured with control cells versus 5 patient samples cultured with CAR-Jurkats. (**D**) Ramos, Raji, HG3 and MEC1 cells were cultured with control or CD19-CAR-Jurkat cells at 1:1 ratio for 48h and then B cells were sorted out. Total cellular CD19 protein expression was determined by Western blotting. GAPDH staining was used as a loading control. (**E**) CD19 mRNA expression from B cells treated as in D was analyzed by quantitative real-time PCR. Mean of two biological replicates is shown (±SD). (**F**) Ramos, Raji, HG3 and MEC1 cells were treated with CD19 antibody Tafasitamab or vehicle alone and CD19 mRNA expression was analyzed by quantitative real-time PCR. Mean of two biological replicates is shown (±SD).

Because the Jurkat cell line is derived from the CD4 lineage, we sought to determine whether this CD19 downregulation was driven exclusively by the CD4+ helper T-cell population. Co-culture with primary CD4+ CAR-T cells yielded significant CD19 loss at both surface and intracellular level mirroring our previous findings **(Supplementary figure 1)**.

Conversely, co-culture with primary CD8+ cytotoxic CAR-T cells induced widespread cell death across multiple evaluated cell lines (Ramos, Mec1, DOHH2, and JVM2). However, the surviving subpopulation of HG3 cells in this condition still exhibited marked CD19 downregulation **(Supplementary figure 2)**. To further pinpoint the effect of CD8+ effectors without the confounding factor of strong cell cytotoxicity, we utilized CD8+ T-cell line HSB-2, which is derived from immature T lymphoblasts and is not cytotoxic. Co-culture with HSB-2 expressing CD19-CAR indeed induced a similar degree of CD19 antigen loss on all the tested B-cell lines, without inducing cell death **(Supplementary figure 3)**.

We next sought to determine the changes at the level of *CD19* mRNA. Quantitative real-time PCR revealed that *CD19* mRNA expression was partially reduced in the majority of tested B-cell lines following co-culture with CAR-Jurkat cells **(Figure 1F)**. To decipher whether the downregulation of CD19 is specific only to CAR-T triggering, we utilized Tafasitamab (TFSM), a humanized cytolytic CD19 antibody. Treating the B-cell lines with Tafasitamab yielded a comparable loss of surface and intracellular CD19 protein **(Supplementary figure 4)**, while only slightly impacting *CD19* mRNA transcripts **(Figure 1G)**. To rule out the possibility of selecting *CD19* mutational variants or clones with genomic deletions through the CAR-T treatment, we performed exon sequencing of the entire *CD19* gene in CAR-T treated cells and revealed a complete absence of any genomic mutations **(Supplementary figure 5)**.

Together, these data demonstrate that unproductive immunotherapy (not immediately resulting in target cell death) mediates profound CD19 loss, which is a non-mutational, post-transcriptional event shared across multiple malignant B-cell lines and primary CLL cells, triggered by both CAR-T and therapeutic antibody modalities.

### Genome-wide CRISPR screening identifies CUL1 and CD81 as key regulators of CD19 antigen expression

In order to understand the molecular mechanisms responsible for CD19 downregulation and to identify genetic factors modulating CD19 expression dynamics, we performed an unbiased, genome-wide CRISPR/Cas9 knockout screen **(Figure 2A)**. Ramos cells were infected with a Brunello knockout library, targeting almost an entire human genome, cells were selected with puromycin and expanded for two weeks. These cells were then co-cultured with primary CAR-T cells at a ratio of 1:5 (Effector:Target) for 48 hours, followed by cell sorting to isolate CD19-high and CD19-low populations. The identity of genes knocked out in these two populations was revealed by next-generation sequencing followed by bioinformatics analysis **(Figure 2B)**. Gene ontology (GO) analysis of the significantly enriched gene hits in CD19-positive population highlighted protein localization pathway as the primary biological process **(Figure 2C)**.

**Figure 2.**
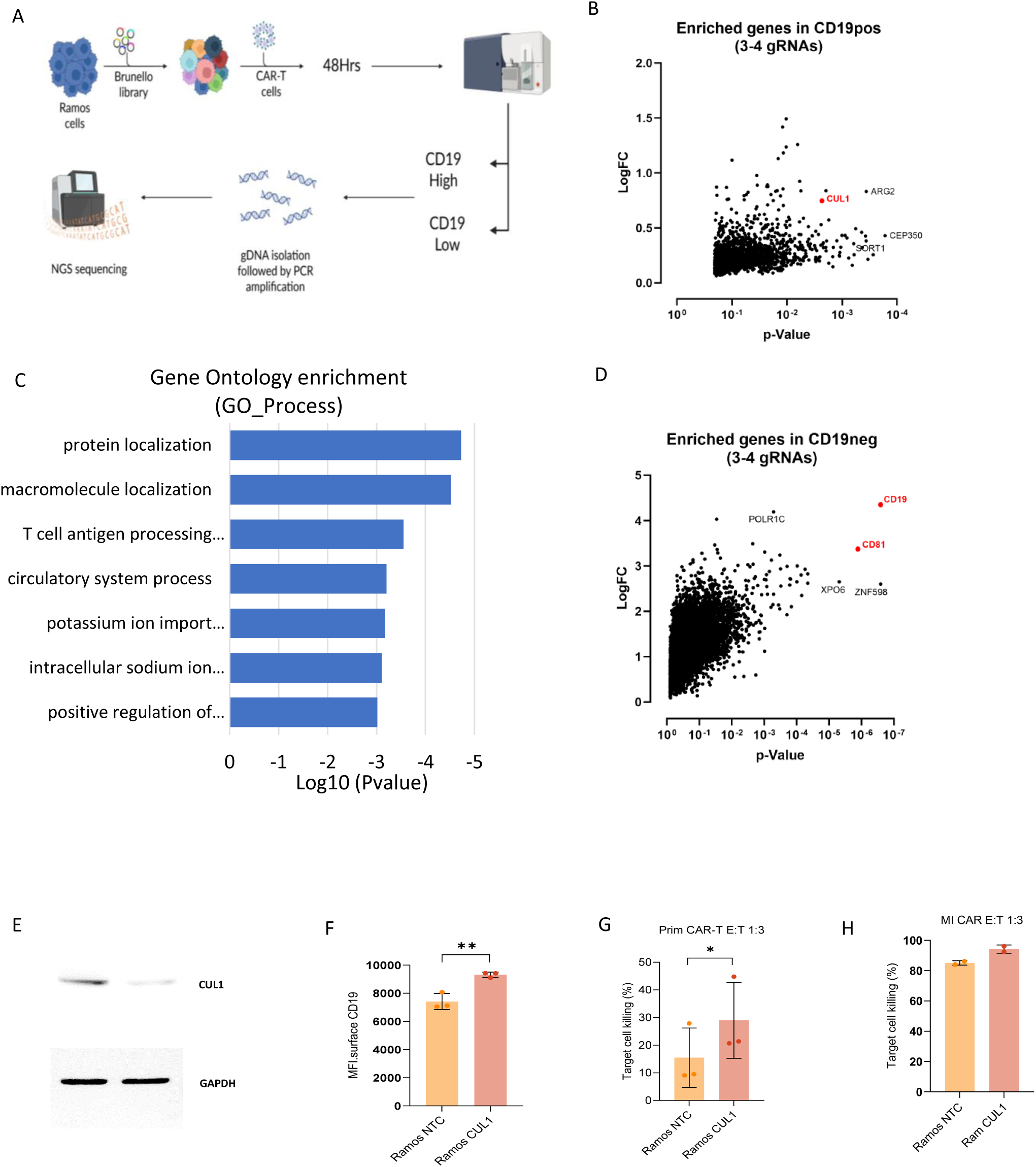
Genome-wide CRISPR knockout screen identifies Cullin-1 and CD81 as the key regulators of CD19 antigen expression. (**A**) A scheme of performed genome-wide CRISPR screen. Brunello CRISPR library was introduced into Ramos B-cell line, cells selected and cultured for 2 weeks. The cells were then mixed with CAR-T cells for 48h and CD19-high and CD19-low B cells were sorted, their genomic DNA isolated and analyzed by NGS. (**B**) Plot depicts -log_10_ p-value and log_2_ fold change of normalized sgRNA counts for the genes, for which at least 3 sgRNAs were found enriched in CD19-positive population. (**C**) Plot as in B displays genes most enriched in the CD19-negative population. (**D**) Gene ontology analysis of the genes found significantly enriched in CD19-positive population (pR0.05). (**E**) Western blot showing Cullin-1 protein expression in Ramos cells transfected with non-targeting (NTC) or Cullin-1-specific (CUL1) sgRNA. GAPDH is used as a loading control. (**F**) CD19 surface expression on Ramos control (NTC) or CUL1-knockout cells was determined by flow cytometry. (**G**) Ramos control and CUL1 knockout cells were mixed with primary CAR-T cells at 1:3 E:T ratio and percentage of killed B cells was determined by flow cytometry via 7-AAD viability staining gated on B cells. (**H**) Ramos control and CUL1-KO cells were mixed with CAR-NK92MI cells at 1:3 E:T ratio and percentage of killed B cells determined as in G.

Notably, our screen identified two genetic regulators of antigen dynamics. Within the CD19-high population, *CUL1* (Cullin-1)—a core scaffolding component of the cullin-RING E3 SCF ubiquitin ligase complex was identified as one of the top enriched hits. Conversely, within the CD19-low population, the tetraspanin chaperone *CD81* emerged as a top candidate **(Figure 2D)**. Importantly, we found *CD19* gene itself as the top hit in the CD19-low population, which served as a positive control and indicated the quality of our screen.

To validate the primary screening hit, we generated *CUL1* knockout (KO) in Ramos cell line, using newly designed specific guideRNAs. Genetic deletion of *CUL1* resulted in a significant baseline increase in surface CD19 expression (Figure 2E, 2F). Consequently, *CUL1*-KO cells exhibited also heightened susceptibility to immune mediated lysis, showing significantly increased killing when challenged with primary CAR-T cells **(Figure 2G)** as well as with another immunotherapeutic modality CAR-NK cells **(Figure 2H)**.

### Pharmacological inhibition of Cullin 1 with pevonedistat prevents CD19 loss and enhances immunotherapeutic efficacy *in vitro*

To translate these genetic findings into a more clinically translatable setting, we mimicked *CUL1* deficiency using pevonedistat, a small molecule inhibitor of the NEDD8-activating enzyme, which is required for cullin neddylation and hence its activation. Pevonedistat treatment induced a rapid increase in surface CD19 expression within 90 minutes and this was sustained for at least 24 hours **(Figure 3A)**. More importantly, pevonedistat effectively rescued CD19 expression on Ramos and MEC1 cells under the selective pressure of CAR-expressing Jurkat T cells **(Figure 3B)**.

**Figure 3.**
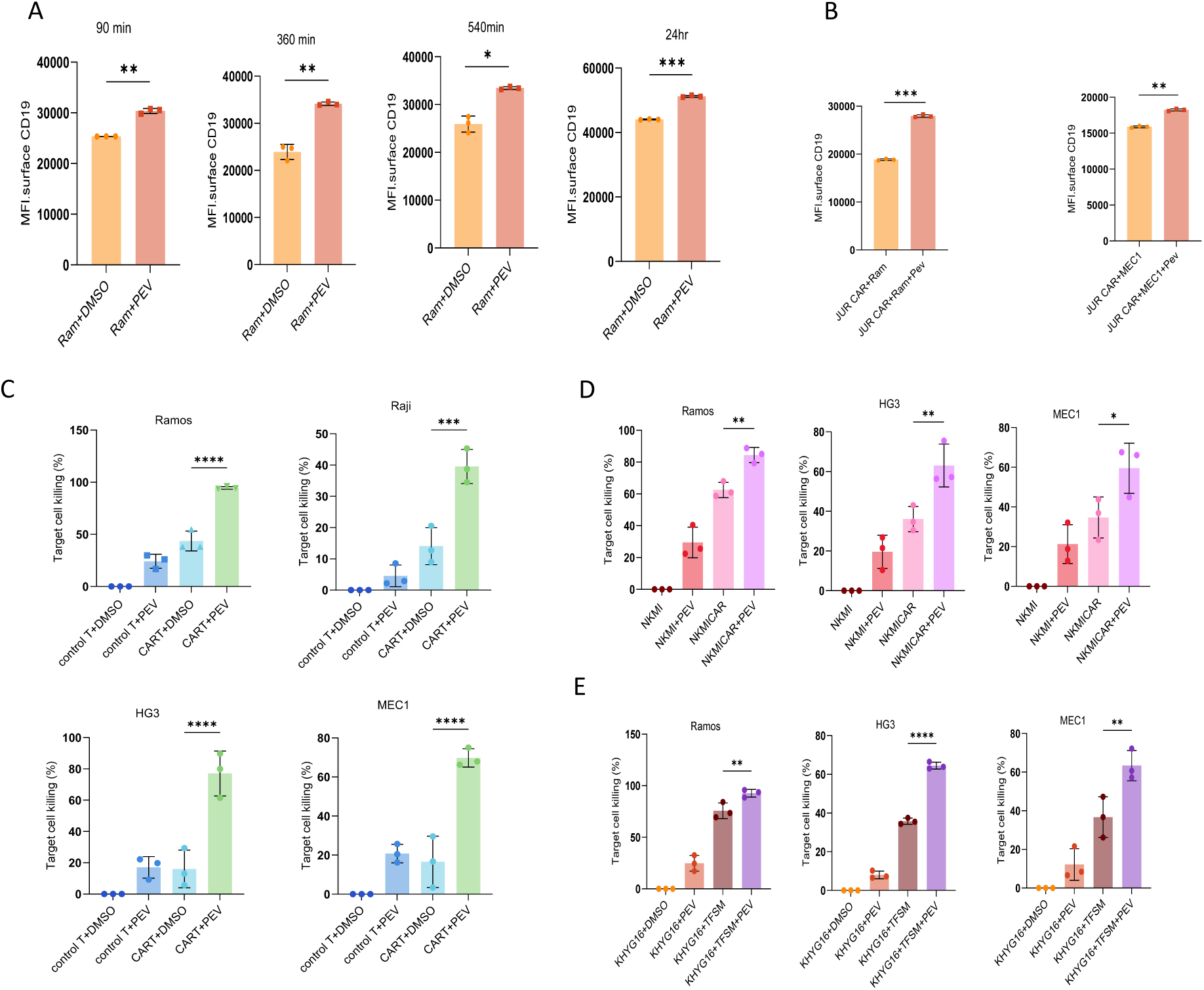
Cullin-1 inhibition prevents CD19 loss and enhances immunotherapeutic killing. (**A**) Ramos cells were treated with Cullin-1 inhibitor pevonedistat or DMSO control for indicated time intervals and the expression of CD19 on the cell surface was determined by flow cytometry. (**B**) Ramos and MEC1 cells were cocultured with CAR-Jurkat cells and concurrently treated with pevonedistat or DMSO control. Surface CD19 expression was determined by flow cytometry. (**C**) Ramos, Raji, HG3 and MEC1 B-cell lines were cocultured with primary control or CAR-T cells together with DMSO or pevonedistat treatment. Percentage of killed B cells after 48h of coculture was determined by flow cytometry measurement of 7-AAD viability staining gated on B cells. (D-E) The same experiment as in C was repeated by culturing Ramos, HG3 and MEC1 cells with control or CAR-NK92MI cells ± pevonedistat (**D**) or by treating them with CD19-targeting antibody tafasitamab plus CD16-expressing KHYG NK cells (KHYG16) ± pevonedistat treatment (**E**).

We next tested whether Pevonedistat-mediated stabilization of CD19 antigen density could potentiate effector cell-mediated cytotoxicity. Indeed, combining Pevonedistat with primary CAR-T cells yielded synergistically strongly enhanced killing against a broad panel of malignant B-cell lines, including Ramos, Raji, HG3, and Mec1 **(Figure 3C)**. Importantly, this sensitization was not specific to just CAR-T cells, but extended to other CD19-targeting effector cell platforms, where pevonedistat significantly boosted CAR-NK mediated cytotoxicity **(Figure 3D)**, as well as antibody-dependent cell cytotoxicity (ADCC) mediated through the combination of CD19-targeting antibody tafasitamab with the FCyRIII (CD16)-expressing NK effector cell line KHYG (KHYG16) **(Figure 3E)**.

### Pevonedistat synergizes with CAR-T, CAR-NK and monoclonal antibodies to enhance killing in primary CLL samples

To confirm the translational relevance of these findings, we evaluated the therapeutic efficacy of Pevonedistat on primary patient samples. Primary CLL patient cells were treated with Pevonedistat in combination with various immunotherapeutic effectors. We observed a significant increase in primary cell lysis when Pevonedistat was combined with primary CAR-T cells **(Figure 4A)**, CAR-NK cells **(Figure 4B)** and tafasitamab-armed KHYG16 cells **(Figure 4C)**. These data confirm that targeting Cullin-1 pathway preserves target antigen availability and maximizes the cytotoxic output of multiple immunotherapy platforms in patient-derived samples.

**Figure 4.**
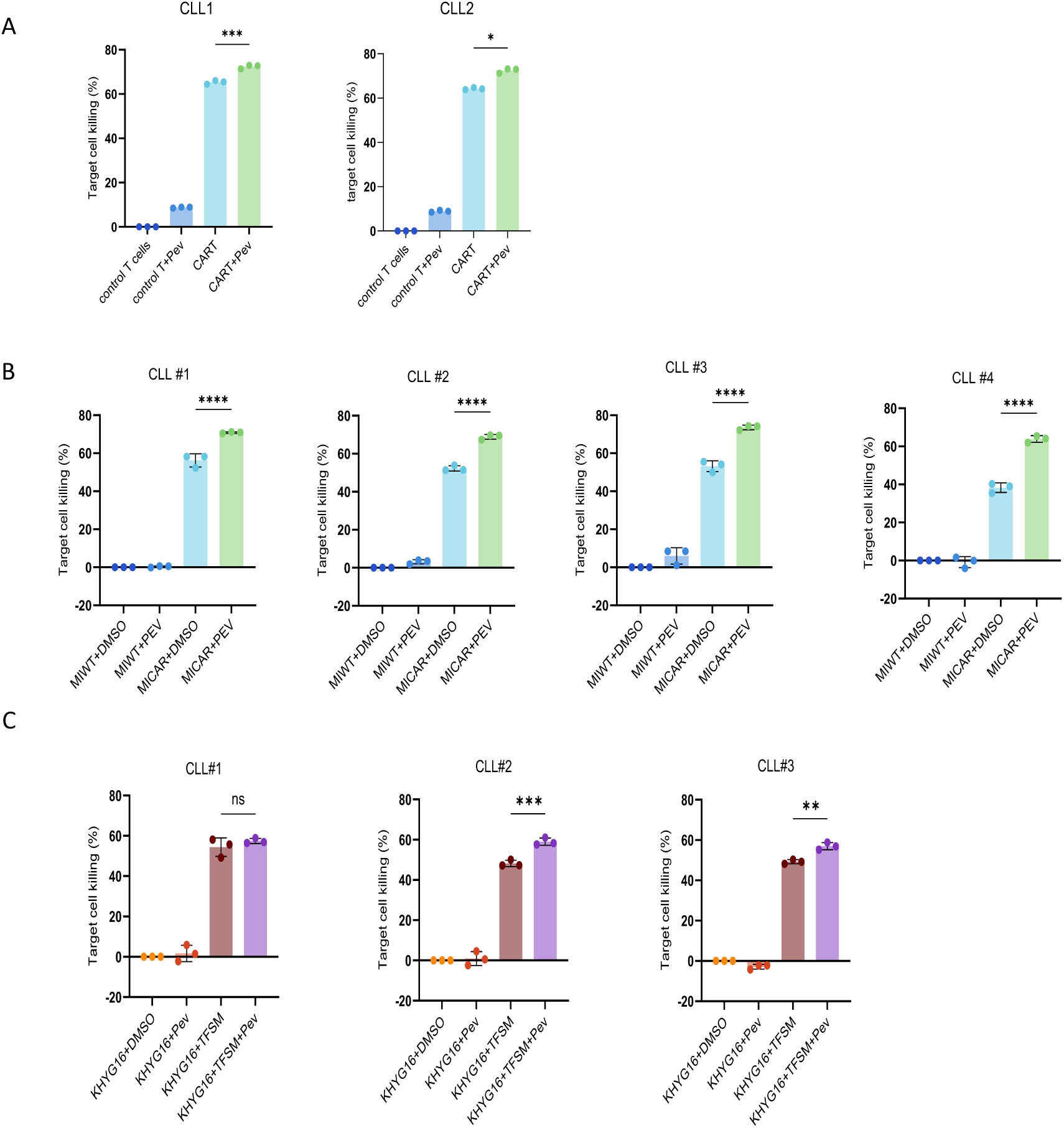
Pevonedistat increases CD19 immunotherapy-mediated killing of primary patient CLL cells *ex vivo.* Primary patient CLL cells were cultured with primary CAR-T cells (**A**), CAR-NK92MI cells (**B**) or with CD19 antibody tafasitamab + CD16-KHYG (**C**) and concurrently treated with pevonedistat or DMSO. Percentage of killed B cells was determined by flow cytometry measurement of 7-AAD viability staining gated on B cells. Three replicates for each patient are shown and mean ± SEM displayed.

### Pevonedistat stabilizes CD19 post-translationally and enhances CAR-T/NK cell activation

We next investigated the molecular mechanism by which Pevonedistat modulates CD19 expression. First, we concluded that the effect is mediated at the protein level, as pevonedistat treatment did not enhance *CD19* mRNA levels **(Figure 5A)**. On contrary, CD19 mRNA seemed to gradually decrease with the time of Pev incubation. Next, we investigated whether protein synthesis is required for the observed upregulation of CD19 protein levels. We therefore treated the cells with protein translation inhibitor cycloheximide (CHX) to block protein synthesis along with Pevonedistat **(Figure 5B)**. We observed the same increase in total CD19 protein expression in both CHX-treated and untreated samples, suggesting that Pev-induced CD19 upregulation does not require synthesis of a new protein. Interestingly, western blot analysis revealed that pevonedistat alone did not directly induce cell intrinsic apoptosis in target B cells, as evidenced by unchanged expression levels of P53, BAD, BAX, and PUMA, alongside a compensatory increase in anti-apoptotic MCL1 **(Supplementary figure 6)**.

**Figure 5.**
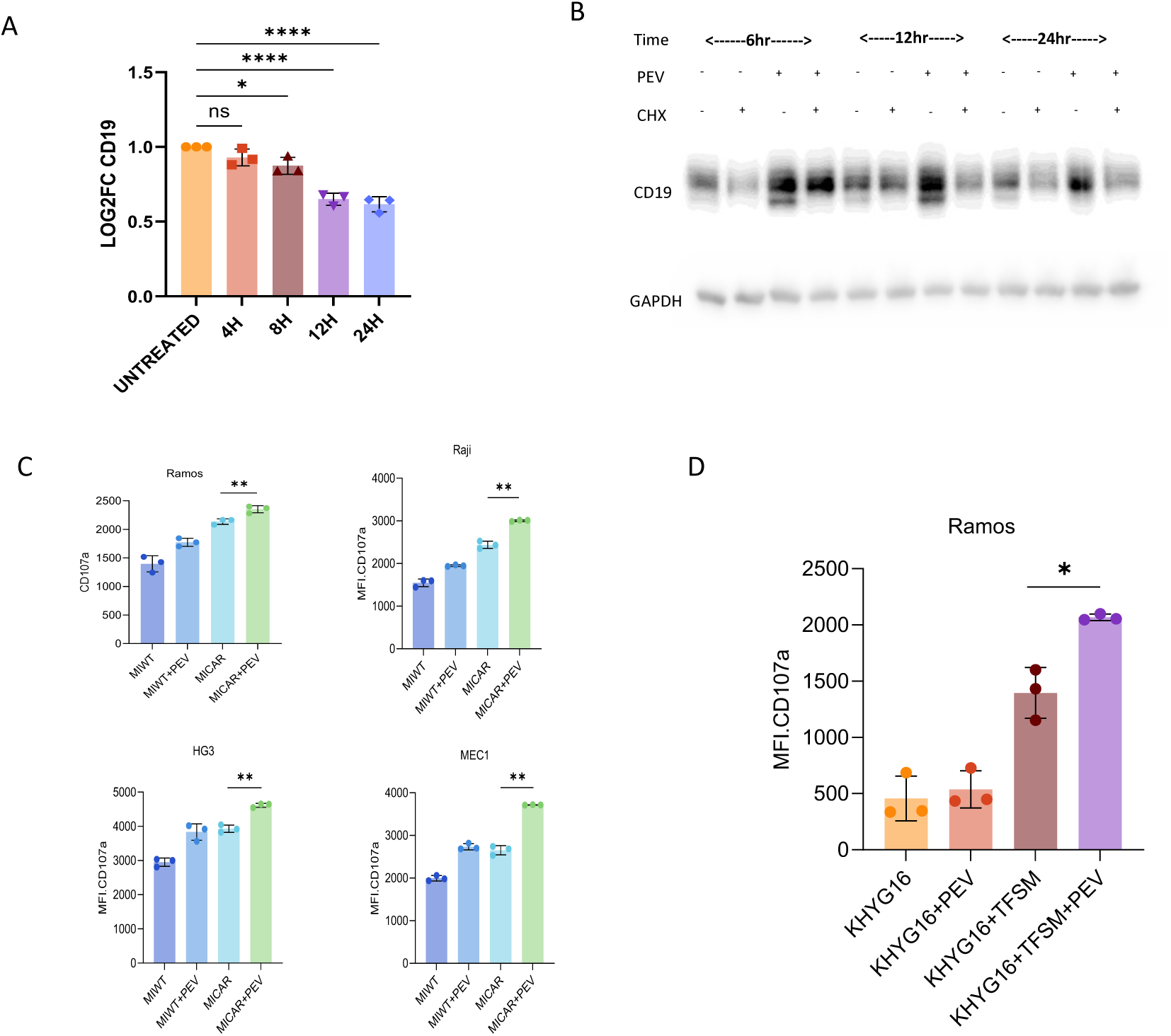
Pevonedistat stabilizes CD19 post-translationally and enhances effector cell activation. (**A**) Ramos cells were treated with pevonedistat for indicated times and CD19 mRNA expression was determined by quantitative real-time PCR. (**B**) Ramos cells were treated with DMSO, cycloheximide (to block protein synthesis), pevonedistat or their combination for indicated times. Total CD19 protein expression was determined by western blotting. GAPDH was used as a loading control. (**C**) Ramos cells were treated with pevonedistat for indicated timepoints and the expression of individual proteins was determined by western blotting. GAPDH was used as a loading control. (**D**) Ramos, Raji, HG3 and MEC1 cells were cultured with CAR-NK92MI cells together with pevonedistat or DMSO control and the expression of CD107a degranulation marker on the surface of NK92MI cells was determined by flow cytometry. (**E**) Ramos cells were cultured with tafasitamab plus CD16-KHYG cells together with pevonedistat or DMSO control and CD107a expression on KHYG16 cells was determined by flow cytometry.

Instead, the enhanced killing was driven by heightened effector cell activation resulting from sustained antigen density. Co-culture of target cells in the presence of pevonedistat led to a substantial upregulation of the degranulation marker CD107a on both CAR-NK and KHYG16 cells (Figure 5C-D), confirming that pevonedistat acts primarily by maintaining a high antigen density required for robust immune synapse formation.

### CUL1 inhibition stabilizes CD19 and drives CD81-dependent immunological synapse formation to potentiate effector cytotoxicity

Finally, as the CD19 stabilization under CUL1 inhibition is not dependent on protein synthesis, we characterized the intracellular trafficking pathway of CD19. To dissect this mechanistically, we treated B-cell lines with a panel of pathway specific inhibitors: dynasore (an endocytosis/dynamin inhibitor), bafilomycin A1 (a lysosomal degradation inhibitor), MG132 (a proteasome inhibitor), or pevonedistat. Accumulation of surface CD19 protein was exclusively observed following treatment with either pevonedistat or dynasore, whereas lysosomal or proteasomal inhibition via bafilomycin or MG132 failed to rescue CD19 levels **(Figure 6A)**. Consequently, functional cytotoxicity assay with the NK-92MI effector system revealed significantly enhanced cell killing upon both dynasore and pevonedistat treatment, nevertheless, pevonedistat-induced increase was even more potentiated compared to dynasore **(Figure 6B)**.

**Figure 6.**
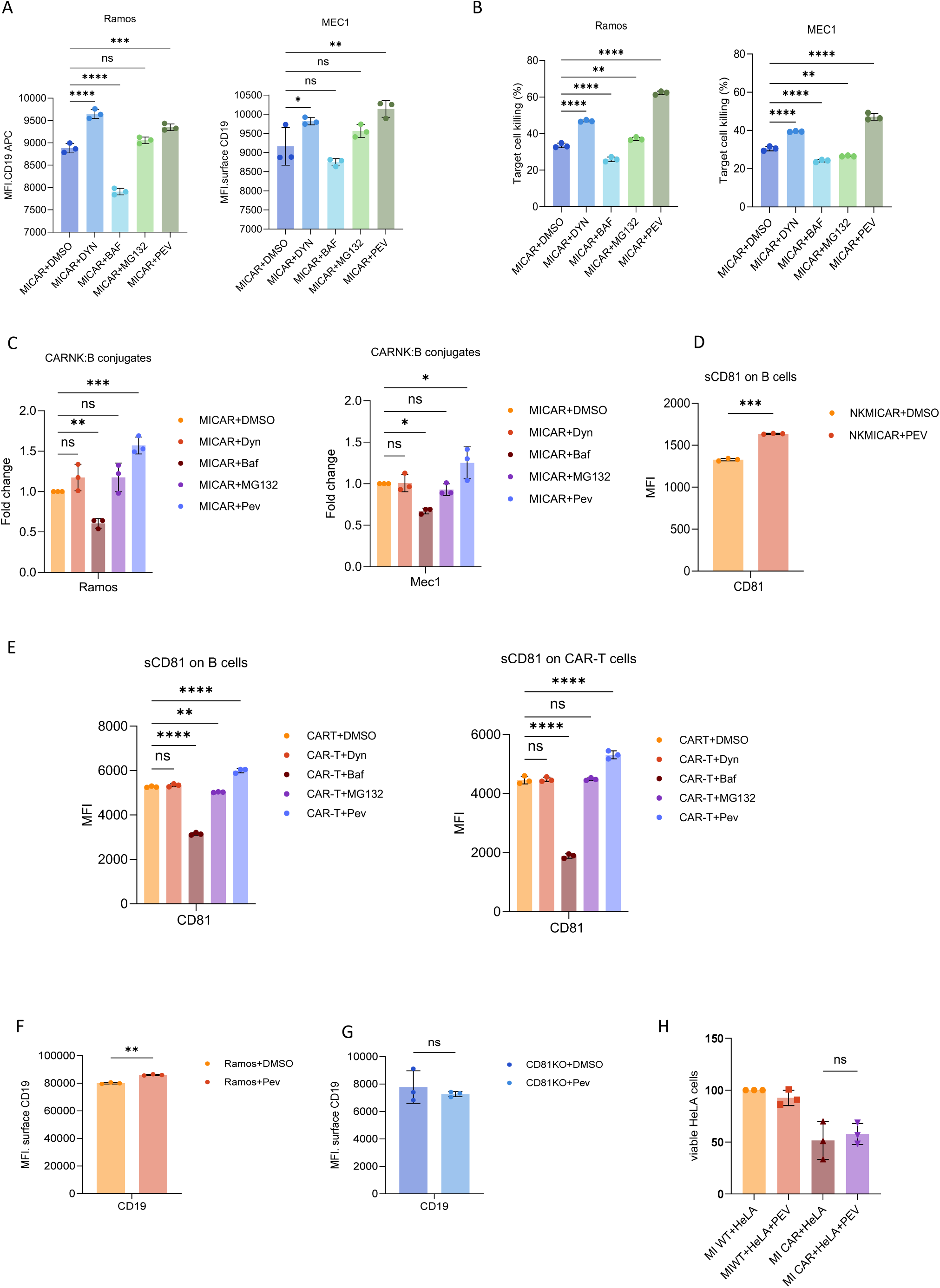
Pevonedistat augments CD19 surface recycling in a CD81-dependent manner. (**A**) Ramos and MEC1 cells were cocultured with CAR-NK92MI cells and treated with DMSO, Dynasore, Bafilomycin, MG132 or Pevonedistat for 48h. CD19 expression levels on the surface of B cells were determined by flow cytometry. (**B**) The experiment as in A was repeated, but, this time, the percentage of killed B cells was determined by 7-AAD staining on gated B cells. (**C**) Ramos cells were cultured with CAR-Jurkats and DMSO or pevonedistat for 48h. Cells were stained with CD19-APC antibody, washed, incubated for 30min and then the extracellular staining was stripped away with the acetic acid. The amount of internalized intracellular stained CD19 protein was measured by flow cytometry. The cells were then incubated for 3h and 16h and the level of recycled surface CD19 staining was determined by flow cytometry. (**D**) Ramos cells were cultured with CAR-Jurkats with DMSO or with pevonedistat and CD81 levels on the surface of Ramos cells (left panel) or Jurkat cells (right panel) were determined by flow cytometry. (**E**) CD81 was knocked out in Ramos cells and the recycling and internalization of CD19 surface protein was assessed in CD81-knockout cells the same way as in C. (**F**) CD19 was exogenously expressed in HeLa cells and they were cocultured with CAR-NK92MI cells with DMSO or with pevonedistat. Percentage of killed HeLa cells was determined by 7-AAD viability staining and gating on HeLa cells. (**G**) Ramos and MEC1 cells were mixed with CAR-NK92MI cells and treated with indicated inhibitors. The percentage of cell-cell conjugates was determined by flow cytometry.

Since pevonedistat increases CD19 surface expression and killing by effector cells, we hypothesized that pevonedistat might impact on the physical interaction between effector and target cells. Indeed, pevonedistat treatment significantly enhanced the formation of stable CAR-NK:B cell conjugates in both Ramos and MEC-1 models, suggesting that pevonedistat increases the immune synapse (IS) formation, a prerequisite for productive cytotoxicity **(Figure 6C)**. Importantly, our initial CRISPR screen identified CD81 as the top enriched hit in CD19-low population. Because CD81 is a critical tetraspanin known to organize membrane microdomains essential for IS formation, we examined whether pevonedistat affects its expression. Flow cytometric analysis revealed that pevonedistat treatment significantly upregulated CD81 expression on both NK effector and target B cells **(Figure 6D)**.

We further validated these findings showing that pevonedistat treatment triggered a significant increase of CD81 expression also on both primary CAR-T cells and HG3 target cells **(Figure 6E)**.

Finally, to confirm that pevonedistat’s effect relies on the CD81 axis, we generated a *CD81*-knockout cell model. In the absence of CD81, cells exhibited a profound loss of CD19, and crucially, pevonedistat treatment completely failed to increase CD19 levels **(Figure 6F)**. To further confirm the necessity of the native CD81 chaperone network, we utilized a HeLa cell system overexpressing CD19. Because HeLa cells lack the B-cell-specific CD81 tetraspanin, the addition of pevonedistat failed to enhance CD19-CAR-NK cell killing against these cells **(Figure 6G)**.

Taken together, our data demonstrate that pevonedistat does not merely prevent passive degradation, but rather actively upregulates CD81 on both effector and target cells to promote immune synapse formation and stabilize surface CD19, thereby potentiating immunotherapeutic efficacy.

## Discussion

The remarkable clinical success of CD19-directed immunotherapies is frequently limited by the emergence of antigen escape. While genetic mutations or alternative splicing account for a subset of relapses^5,6^, the majority of resistant tumors retain intact CD19 transcripts despite a profound loss of surface protein^6^. In this study, we demonstrated that immunotherapeutic selective pressure rapidly induces a non-mutational, post-translational downregulation of CD19 in response to both CAR-T/NK and anti-CD19 monoclonal antibody platforms. Through an unbiased genome-wide CRISPR knockout screen, we identified the core component of E3 ubiquitin ligase complex Cullin-1 and the tetraspanin chaperone CD81 as central regulators of this CD19 antigen escape mechanism.

Cullin-1 is an essential scaffold protein of a Cullin-RING-E3 ubiquitin ligase complex (CRL), also called SCF complex (SKP1-CUL1-F-box protein), which is responsible for ubiquitination of multiple target proteins involved in cell cycle progression, signal transduction and transcription^13^. Our findings functionally validate the CRL network as a critical, druggable vulnerability in the maintenance of tumor antigenicity towards immunotherapy agents. We demonstrated that pharmacological inhibition of CRLs using pevonedistat successfully stabilizes CD19 surface expression under immunotherapeutic pressure. Crucially, this intervention synergistically enhances the cytotoxicity of primary CAR-T cells, CAR-NK cells, and tafasitamab+CD16-NK cells combination against a broad panel of malignant B-cell lines and primary patient-derived CLL samples.

Mechanistically, our data reveal that the ubiquitination pathways governing CD19 trafficking do not control the rate of its endocytosis. Instead, CUL1 inhibition by pevonedistat prevents CD19 degradation and accelerates the surface recycling of internalized CD19. We show that this process is absolutely dependent on the upregulation and interaction with the CD81 chaperone. In the absence of an intact CD19/CD81 complex, such as in our CD81-knockout B cells or in CD81-deficient HeLa cell model, pevonedistat fails to induce CD19 surface recycling, resulting in aberrant intracellular accumulation of the antigen. This aligns with previous structural studies highlighting CD81’s obligate role in exporting CD19 to the plasma membrane and organizing it into tetraspanin-enriched microdomains^22^. The stabilization of these membrane microdomains is paramount for immunotherapeutic efficacy. Indeed, it was shown that CAR-T effector cells require a higher antigen density threshold as compared to classical T-cell receptors in order to form a stable immunological synapse, become fully activated, and trigger robust degranulation and cytokine production^23–25^. By coordinating CD81 upregulation, preventing the proteasomal degradation of CD19 and increasing its cell surface recycling, pevonedistat significantly augments the dynamics of effector – target cell (T - B cell) conjugate formation. This sustains the structural integrity of the immune synapse, maximizing the cytotoxic output of the effector cells.

Interestingly, beside CD81, the top hits revealed in our CRISPR screen that were required for proper CD19 surface maintenance included SORT1 and CEP450. Sortilin (SORT1) functions as a master sorting receptor governing Golgi to endosome trafficking, while the centrosomal protein CEP450 coordinates vesicle tethering and microtubule dependent transport to the plasma membrane (*Ref*.). These findings further strongly support the model where antigen escape is driven by a highly coordinated tumor-intrinsic vesicular trafficking network tightly adjusting antigen levels.

Recent studies have highlighted trogocytosis as a driver of CD19 loss^26,27^. Nevertheless, in our co-cultures, we observed rapid loss of surface CD19 in B cells, but did not detect any CD19 acquisition on the effector cells (data not shown). Therefore, tumor-intrinsic endocytic trafficking constitutes a distinct and potent mechanism of antigen escape that operates independently of trogocytosis-mediated antigen extraction. By accelerating the recycling of this internal pool, pevonedistat directly neutralizes this intrinsic evasion pathway.

The requirement for simultaneous administration of pevonedistat during effector-target engagement perfectly supports the highly dynamic trafficking kinetics of the CD19/CD81 co-receptor complex. Our data demonstrate that pre-treating tumor cells with the CRL inhibitor is insufficient to sustain antigen expression. This indicates that aberrant CD19 endocytosis is not a high turnover background process, but rather a rapid onset event explicitly triggered by the formation of the immunological synapse. For pevonedistat to effectively rescue surface expression, it must be present at the moment of CAR-encounter and its triggering of CD19 antigen internalization. By simultaneously applying selective immunotherapeutic pressure and CRL inhibition, the targeted cells may be forced into an internal trafficking loop: as the CAR-T synapse drives the endocytosis of the CD19/CD81 complex, the concurrent blockade of CUL1-SCF ubiquitination immediately intercepts this trafficking, diverting the intact complex through the recycling pathway back to the plasma membrane. This real time kinetic coupling is essential to sustain the high density antigen microdomains required for maximal cytotoxic output. Importantly, our findings were valid in multiple cell lines with variable CD19 antigen levels, having low, medium or high CD19 expression.

The translational implications of these findings are substantial. Because pevonedistat has already undergone extensive Phase I and Phase II clinical evaluation with a well-characterized safety profile^28^, our data provide a strong mechanistic rationale for its repurposing as a combinatorial agent. Administering CRL inhibitors concurrently with adoptive cell therapies or monoclonal antibodies could minimize the onset of antigen-low escape scenarios, deepening initial responses and prolonging remissions and thereby improving progression-free survival in patients with relapsed or refractory B-cell malignancies.

## Supporting information

Supplementary figures with legends

## Acknowledgements

The authors would like to thank Dr. Sarka Pavlova for providing the primary patient CLL samples, Thesis Advisory Committee members Dr. Michal Simicek and Dr. Jamal Alzubi for their helpful suggestions, and all the members of Functional Genomics research group for insightful discussions and help with experimental procedures.

We acknowledge the Core Facility Genomics and the Core Facility Bioinformatics (both at CEITEC MU) supported by the NCMG research infrastructure (LM2023067 funded by MEYS CR), and Flow cytometry laboratory at CEITEC MU supported by the EATRIS-CZ research infrastructure (LM2023053 funded by MEYS CR) for their support with obtaining scientific data presented in this paper. This study was funded by the research grant from the Czech Science Foundation (project no. 22-35273S), the project National Institute for Cancer Research (Program EXCELES, ID Project No. LX22NPO5102) - Funded by the European Union – Next Generation EU, and project MUNI/A/1733/2025.

## Authorship contributions

N.V.G. designed and performed the experiments, analyzed and interpreted the data, wrote the manuscript; T.L. helped with data collection, data analysis and interpretation; M.S. designed the experiments, analyzed and interpreted the data, wrote the manuscript.

