## Supplementary figures with legends for "Genome-wide CRISPR Screening Reveals Cullin-1 as a Therapeutic Target Enhancing Efficacy of CD19-directed Immunotherapy"

S1A

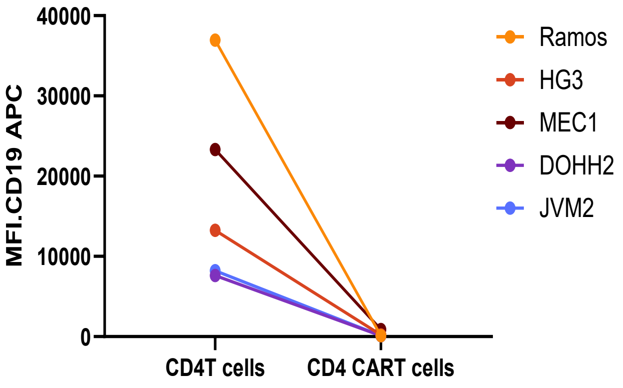


S1B

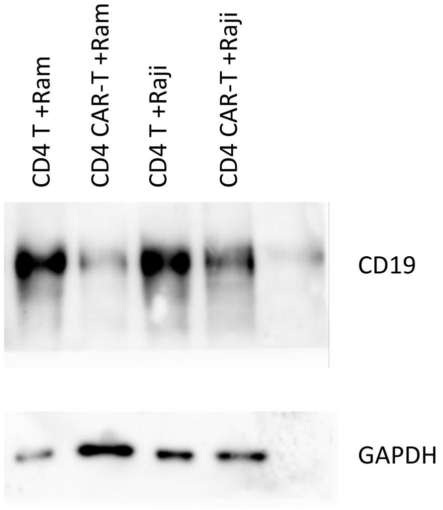


**Figure S1. Surface and total CD19 expression in B cell lines following co-culture with CD4+ CAR T cells. (A)** Flow cytometry analysis of surface CD19 expression. A panel of Azurite-expressing B cell lines (RAMOS, HG3, MEC1, DOHH2, and JVM1) were co-cultured with either primary CD4+ T cells (control) or CD4+ CAR T cells at a 1:3 ratio for 48 hours. **(B)** Western blot analysis of total CD19 protein levels. Azurite-expressing RAMOS and Raji cells were co-cultured under identical conditions for 48 hours, then isolated via fluorescence-activated cell sorting (FACS). Whole-cell lysates from the sorted B cells were probed for CD19, with GAPDH serving as the loading control.

S2

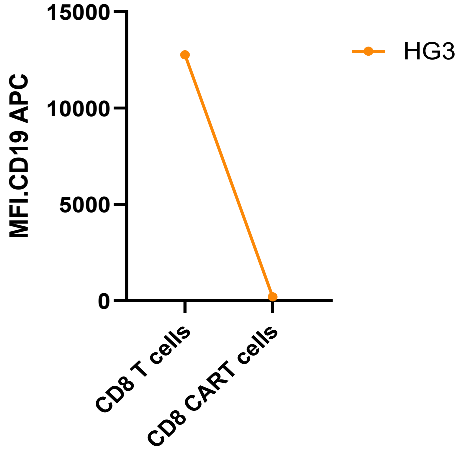


**Figure S2. Surface CD19 expression on HG3 cells following co-culture with CD8+ CAR T cells.** Flow cytometry analysis quantifying the Mean Fluorescence Intensity (MFI) of surface CD19. Azurite-expressing HG3 cells were co-cultured with either primary CD8+ T cells (control) or CD8+ CAR T cells at a 1:3 ratio for 48 hours. Following the co-culture, CD19 MFI was specifically evaluated on the viable, Azurite-positive cell population.

S3

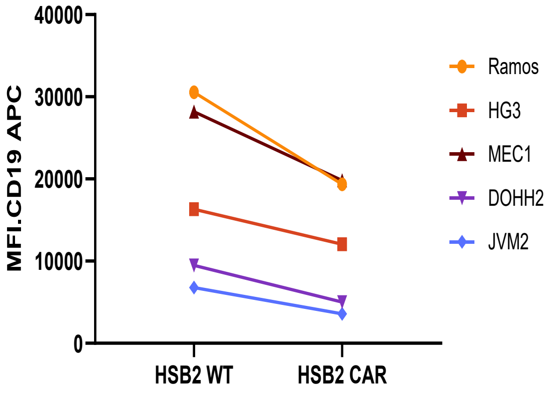


**Figure S3. Surface CD19 expression in B cell lines following co-culture with CD8+ HSB-2 CAR cells.** Flow cytometry analysis of surface CD19 expression. A panel of Azurite-expressing B cell lines (RAMOS, HG3, MEC1, DOHH2, and JVM1) were co-cultured with HSB-2WT cells or HSB-2 CAR cells at a 1:3 ratio for 48 hours. Surface CD19 levels were measured on the viable, Azurite-positive B cell populations

S4A

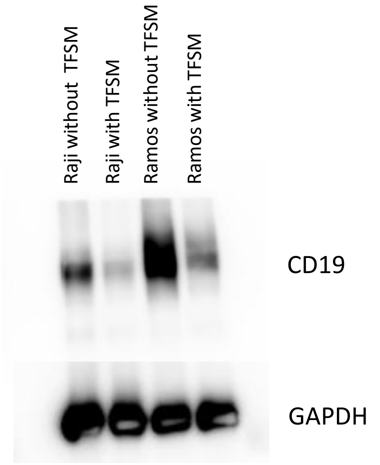


S4B

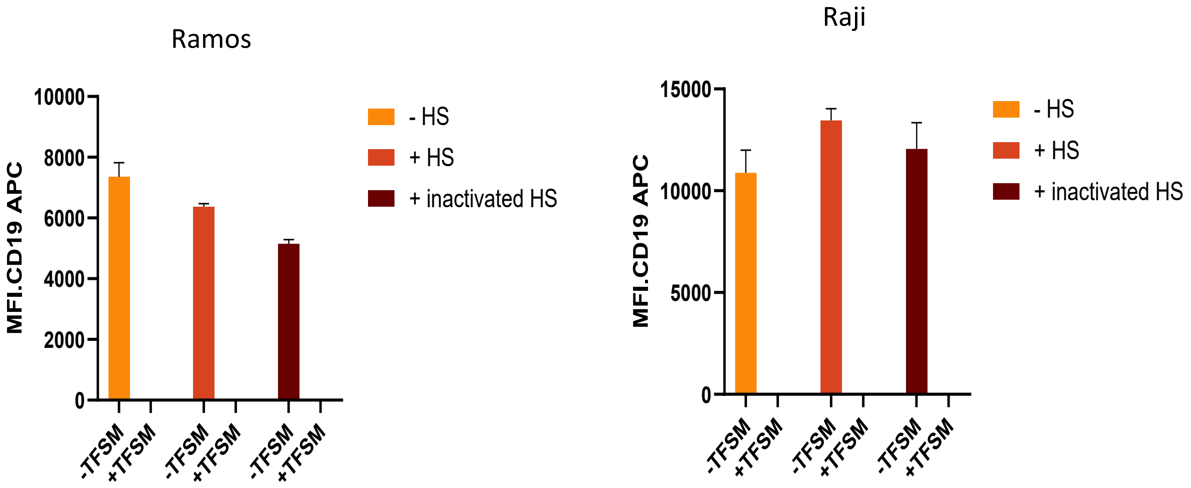


**Figure S4. Effect of tafasitamab and human serum on CD19 expression in Ramos and Raji cells.** **(A)** Western blot analysis of total CD19 protein levels. Ramos and Raji cells were cultured in the presence or absence of tafasitamab for 48 hours. Following treatment, whole-cell extracts were probed for CD19, with GAPDH serving as the loading control. **(B)** Flow cytometry analysis evaluating surface CD19 levels. Cells were treated for 24 hours under three conditions: tafasitamab alone (without serum), tafasitamab with active human serum, or tafasitamab with heat-inactivated human serum. Following incubation, surface CD19 expression was measured.

S5

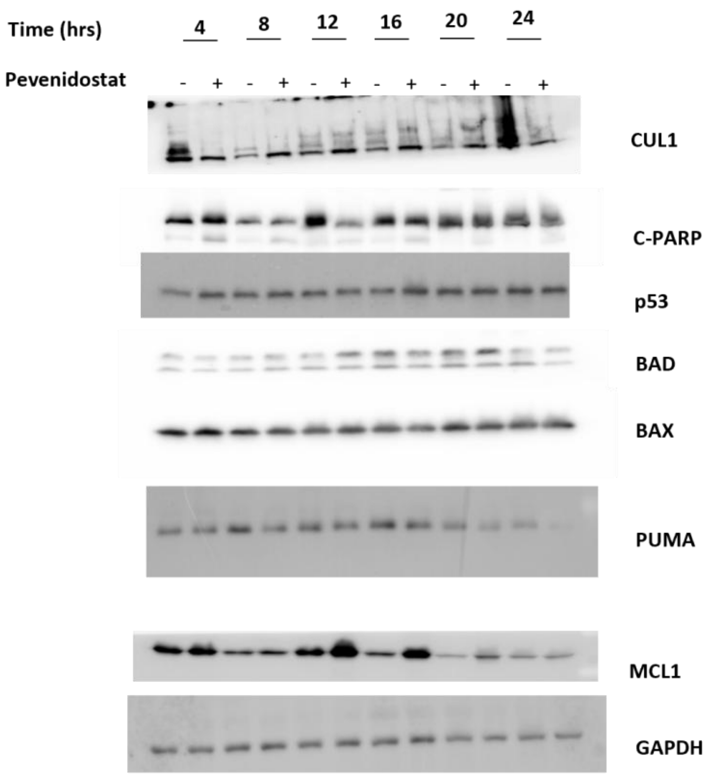


**Figure S5. Time-course analysis of apoptotic markers following pevonedistat treatment.** Western blot analysis of whole cell lysates. Ramos cells were treated with either pevonedistat or DMSO (vehicle control). Cells were harvested every 4 hours over a 24-hour period (4, 8, 12, 16, 20, and 24 hours post-treatment). Lysates were probed for CUL1, cleaved PARP (c-PARP), p53, BAD, BAX, PUMA, and MCL1 to evaluate the induction of apoptosis. GAPDH was used as the loading control across all time points

S6

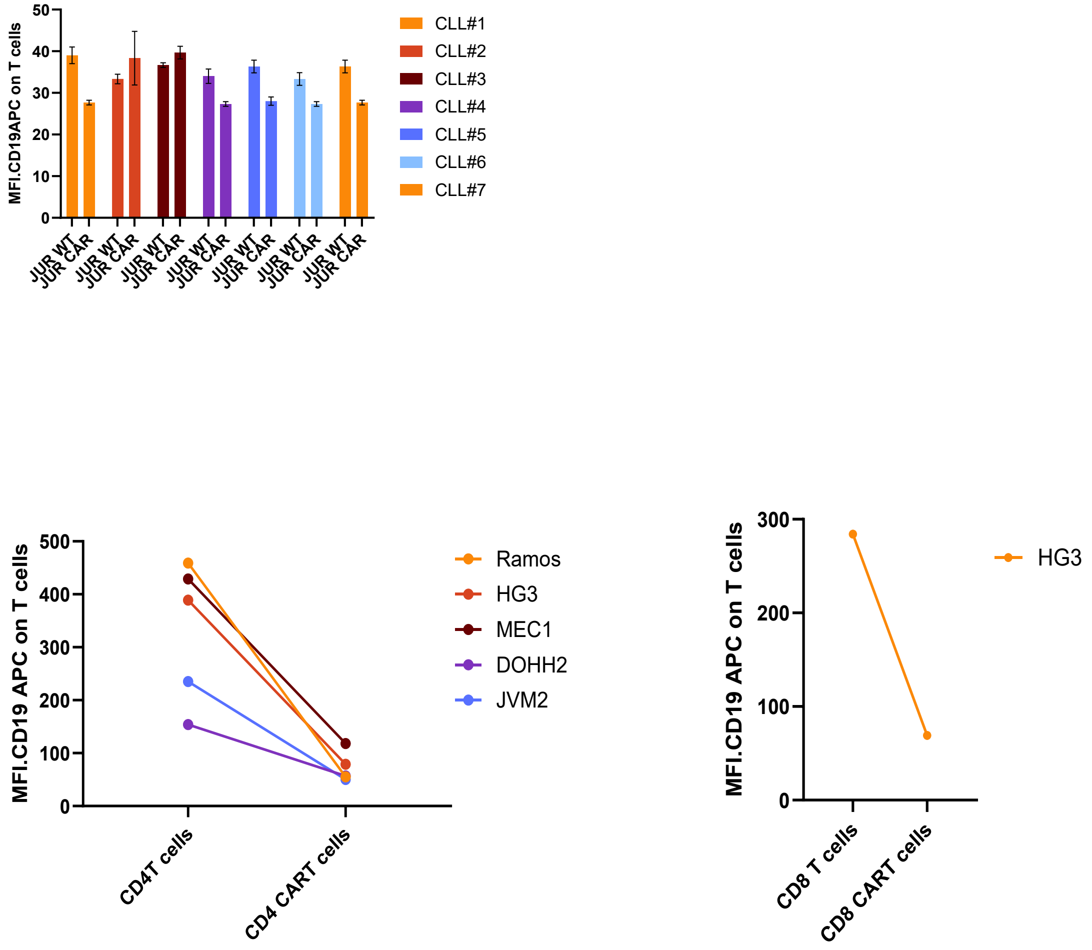


**Figure S6. Evaluation of CD19 trogocytosis by effector T cells and CAR cells.** Flow cytometry analysis assessing the transfer of surface CD19 from target B cells to effector cells via trogocytosis. Data correspond to the 48-hour co-culture experiments detailed in Figure 1C, Figure S1A, and Figure S3. To evaluate trogocytosis, flow cytometry data were re-analyzed by specifically gating on the effector cell populations (primary CD4+ T cells, primary CD8+ T cells, and Jurkat cells) rather than the B cells. The acquisition of CD19 by the effector cells was quantified by measuring the CD19-APC signal within these gated populations.
