## Supplementary table for "Genome-wide CRISPR Screening Reveals Cullin-1 as a Therapeutic Target Enhancing Efficacy of CD19-directed Immunotherapy"

| Product | company | cat. number |
| --- | --- | --- |
| CD19APC | Sony Biosciences | 2111060 |
| CD19FITC | Sony Biosciences | 2111030 |
| CD3 APC | Sony Biosciences | 2186590 |
| CD20FITC | Sony Biosciences | 2111520 |
| CD81 APC | Sony Biosciences | 2347550 |
| LAMP1 APC | Sony Biosciences | 2243095 |
| CD21 PE | Sony Biosciences | 2374515 |
| CD3 FITC | Sony Biosciences | 2186530 |
| CD8a Monoclonal Antibody (SK1), PE-Cyanine7, eBioscience™ | ThermoFisher Scientific | 25-0087-41 |
| APC anti-human CD21 | Sony Biosciences | 2374525 |
| Rabbit Anti-Mouse FMC63 scFv Monoclonal Antibody, Alexa Fluo | Bioswan | 200102 |
| 7AAD | ThermoFisher Scientific | A1310 |
| Sytox blue (dead cell stain for FCM) | ThermoFisher Scientific | S34857 |
| CD19 (Intracellular Domain) (D4V4B) XP® Rabbit mAb | Cell Signaling Technology | 90176S |
| Pro-Survival Bcl-2 Family Antibody Sampler Kit II | Cell Signaling Technology | 17229T |
| Pro-Apoptosis Bcl-2 Family Antibody Sampler Kit II | Cell Signaling Technology | 98322T |
| CUL1 Antibody | Cell Signaling Technology | 4995S |
